# A precursor of reading ability? 3-month-old infants easily learn to pair a phoneme with a visual shape

**DOI:** 10.1101/466318

**Authors:** Karima Mersad, Claire Kabdebon, Ghislaine Dehaene-Lambertz

**Author notes:** **Corresponding author:** Karima Mersad, Laboratoire Vision Action Cognition, 71 avenue Edouard Vaillant, 92774 Boulogne Billancourt Cedex, France.

## Abstract

Being aware of the phonemes that compose syllables is difficult without having learned to read an alphabetic script, a skill generally acquired around 5 to 7 years of age. Nevertheless, preverbal infants are particularly good at discriminating syllables that differ by a single phoneme. Do they perceive syllables as a whole unit or can they become aware of the underlying phonemes if their attention is attracted to the relevant level of analysis? We trained 3-month-old infants to pair two consonants, co-articulated with different vowels, with two visual shapes. Using event-related potentials, we show that infants generalize the learned associations to new syllables with respect to a familiarization phase. The systematic pairing of a visual label with a phonetic category is thus easy to learn, suggesting that the main process underlying reading (i.e., grapheme-phoneme pairing) is grounded in the early faculties of the human linguistic system.

## Introduction

Two words with very different meanings can differ by only a single phoneme. Despite important acoustic variations created by overlapping speech gestures, the emotional state of the speaker, or vocal idiosyncrasies, humans categorically discriminate syllables that vary by a single phoneme. Human infants are also capable of subtle phonemic discrimination (*1, 2*) and can easily account for acoustic variations in co-articulation to correctly categorize, for example, the syllables /bi/ and /bo/ as being closer to /ba/ than /ga/ (*3, 4*). Using event-related potentials (ERPs), we recently replicated these behavioural results in 3-month-old infants: a change in the first consonant of CV syllables evoked a mismatch response despite different co-articulated vowels (*5*). However, this observation might only indicate that in an acoustic-phonetic space, syllables that share more features are perceptually closer than those that share less features (*6*) rather than infants having explicit knowledge of phonetic categories. Indeed, even though illiterate adults can hear the difference between syllables, they have no awareness about their elementary constituents and cannot easily decompose them into their phonetic segments nor easily exchange phonemes between syllables (eg., convert John Lennon into Lohn Jennon).

Hence, the explicit representation of speech as a sequence of phonetic units does not arise spontaneously during development but from specific training in an alphabetic system (*6*). During this process, an implicit phonetic representation is converted into an explicit one by re-coding phonemes into visual symbols. As stated by Karmiloff Smith (*7*), this method of acquiring knowledge is specifically human: information already present in the system is redescribed in a different representational format, rendering it more explicit and manipulable.

Here, we asked whether infants might be able to go beyond implicit representations of phonemes and access sub-syllabic components if we attracted their attention to such features, similar to when older children learn to read. Several experiments have reported that providing a steady label for different exemplars of one category helps infants form a unified representation of the category of which they would have otherwise remained unaware. For example, naming different pictures of dinosaurs with the same non-word, or even with lemur vocalization, helps 3-month-olds differentiate pictures in this category from those of fishes (*8*). Similarly, associating two objects with two different phonetic categories helps 9-month-olds perceive a difficult non-native contrast (*9*). We recently proposed that human infants benefit from symbolic representations (*10*) and spontaneously infer that if a common arbitrary sign is associated with a set of stimuli, they should search for shared features among the stimuli, hence directing their attention to the relevant features of analysis.

To this end, we trained twenty 3-month-old infants to associate a specific phoneme with a specific visual shape (figure 1). During a familiarization phase (30 trials), a syllable /bX/ (or /gX/), with a vowel X randomly drawn from the set 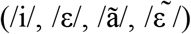, was presented, followed 1.5 s later by a yellow “+” image (or a red “%”). During the subsequent test phase, we examined the generalization of the association by adding two novel syllables /ba/ and /ga/, which were associated with both shapes with equal probability. We reasoned that if infants represent syllables as a whole, without access to the constitutive phonemes, they should have no expectation for a specific shape after hearing the new syllables /ba/ and /ga/. On the contrary, if they understand the relation between the visual shape and the common consonant, they should exhibit a surprise response from an incongruent pairing even in the context of a novel vowel, revealing that they can represent sub-syllable constituents.

**Fig.1.**
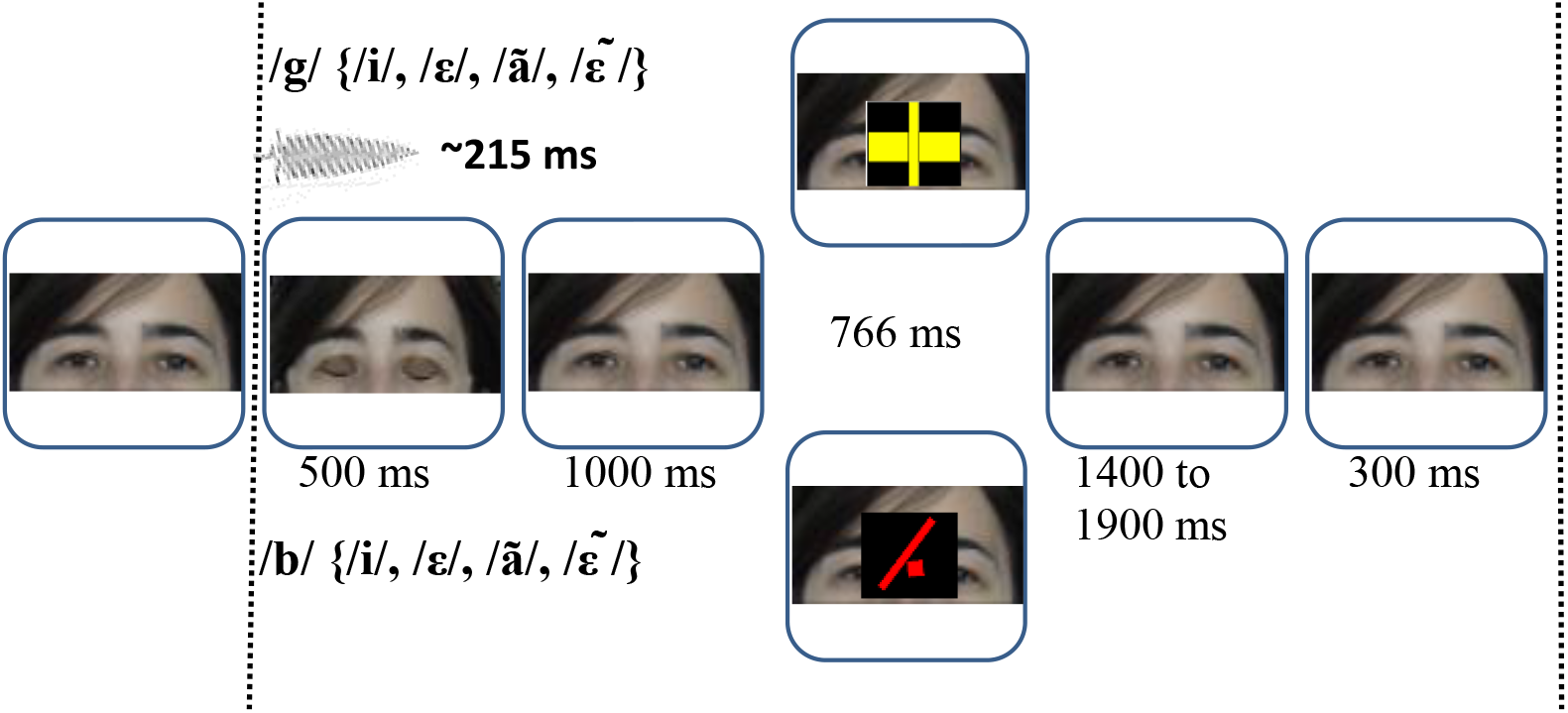
Trial structure. A syllable /bx/, or /gx/, with x randomly chosen from 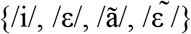 is synchronously presented with blinking eyes, followed by a geometrical shape presented over the eyes 1500 ms after the syllable onset. One consonant is associated with one shape (e.g., /g/ with the yellow “+” and /b/ with the red “%”), counterbalanced across infants. During the test phase, two new syllables /ba/ and /ga/ were added to the set, associated with either of the two shapes with a fifty-fifty probability. The correct vs. incorrect association between the shape and the consonant defines congruent vs. incongruent trials. Incongruency could only be perceived if infants were able to notice the constant association between a consonant and a shape.

We focused our analyses on the P1 and P400, the classical visual ERP components at this age. We considered the responses evoked not only by the onset of the associated shape but also its offset, which was immediately followed by the reappearance of a picture of a woman’s eyes (our fixation attractor, see figure 1) because eyes robustly elicit evoked responses in infants (*11, 12*). In similar paradigms using sudden incongruency in attentive infants, two distinct processing stages have been reported: an early ERP component which reflects a prediction error signal in sensory systems and a late component resulting from an orientation of attention to the unexpected event. The later component is particularly slow in infants, around 1 second (*5, 13–18*) and may extend over subsequent events, reducing the amplitude of usual components. Hence, in our paradigm, the observed effect should be a reduction in the responses to the reappearance of the eyes.

## Results

We compared the visual evoked potentials (ERPs) evoked by congruent and incongruent shapes following two novel syllables /ba/ and /ga/. Trials were first collapsed across congruent and incongruent conditions to identify the P1 and the P400 (figure 2) corresponding to the onset and offset of the shape, over a cluster of occipital and temporal electrodes as commonly performed in the literature to study visual ERPs (*16, 17*). Then, the voltage averaged over the time-window surrounding the peak of each component and over the cluster of electrodes in each subject entered a t-test to investigate differences between congruent and incongruent trials.

**Fig. 2.**
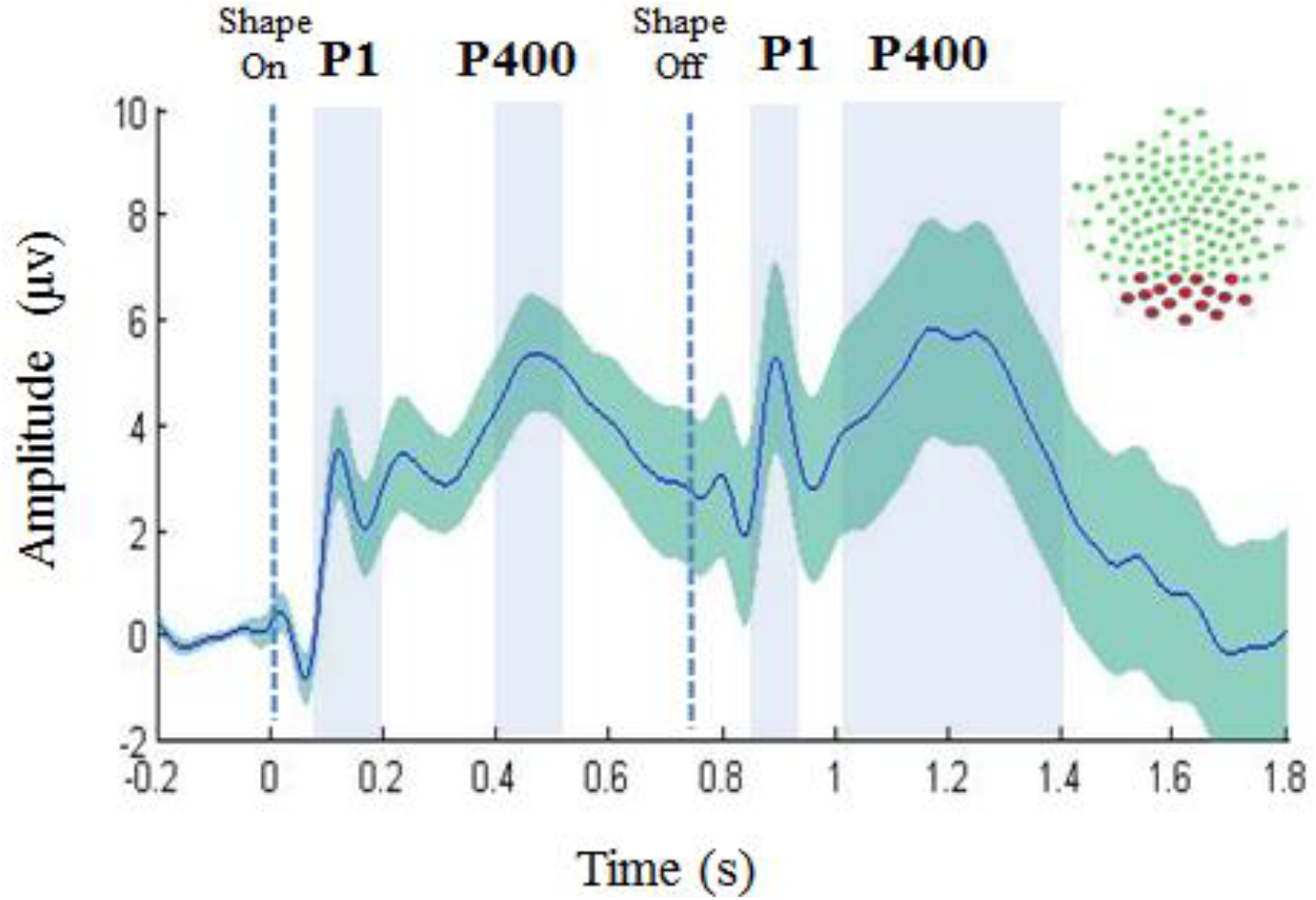
ERP components in response to shape. Grand-average of all trials recorded from the occipito-temporal channels (red dots on the net topography). Blue rectangles indicate the time-windows of the visual ERP components, P1 (100 to 200 ms) and P400 (400 to 500 ms), first after the onset of the shape and, second, P1 (100 to 200 ms) and P400 (250 to 650 ms) after the offset of the shape and reappearance of the eyes. Times-windows were centred on the maximum of the peak amplitude.

A difference of voltage amplitude between conditions slowly built up (figure 3) to significance after the disappearance of the shape, during the P1 (100 to 200 ms) and P400 (250 to 650 ms) components evoked by the reappearance of the eyes (P1: t(19) = 2.88, p = .009, congruent trials: M = 8.28 μv, SE = 3.73 vs. incongruent trials M = −1.68 μv, SE = 4.10; effect size = .64; P400: t(19) = 3.49, p = .002, congruent trials: M = 12.51μv, SE = 4.04 vs. incongruent trials M = -.09 μv, SE = 3.49; effect size = .69). By contrast, we did not observe any sensory preparation for the shape. Precisely, neither the P1 (100 to 200 ms) nor the P400 (400 to 500 ms) component elicited by the shape onset was modulated by the experimental condition (P1: t < 1; P400: t (19) = 1.54, p =.13).

**Fig. 3.**
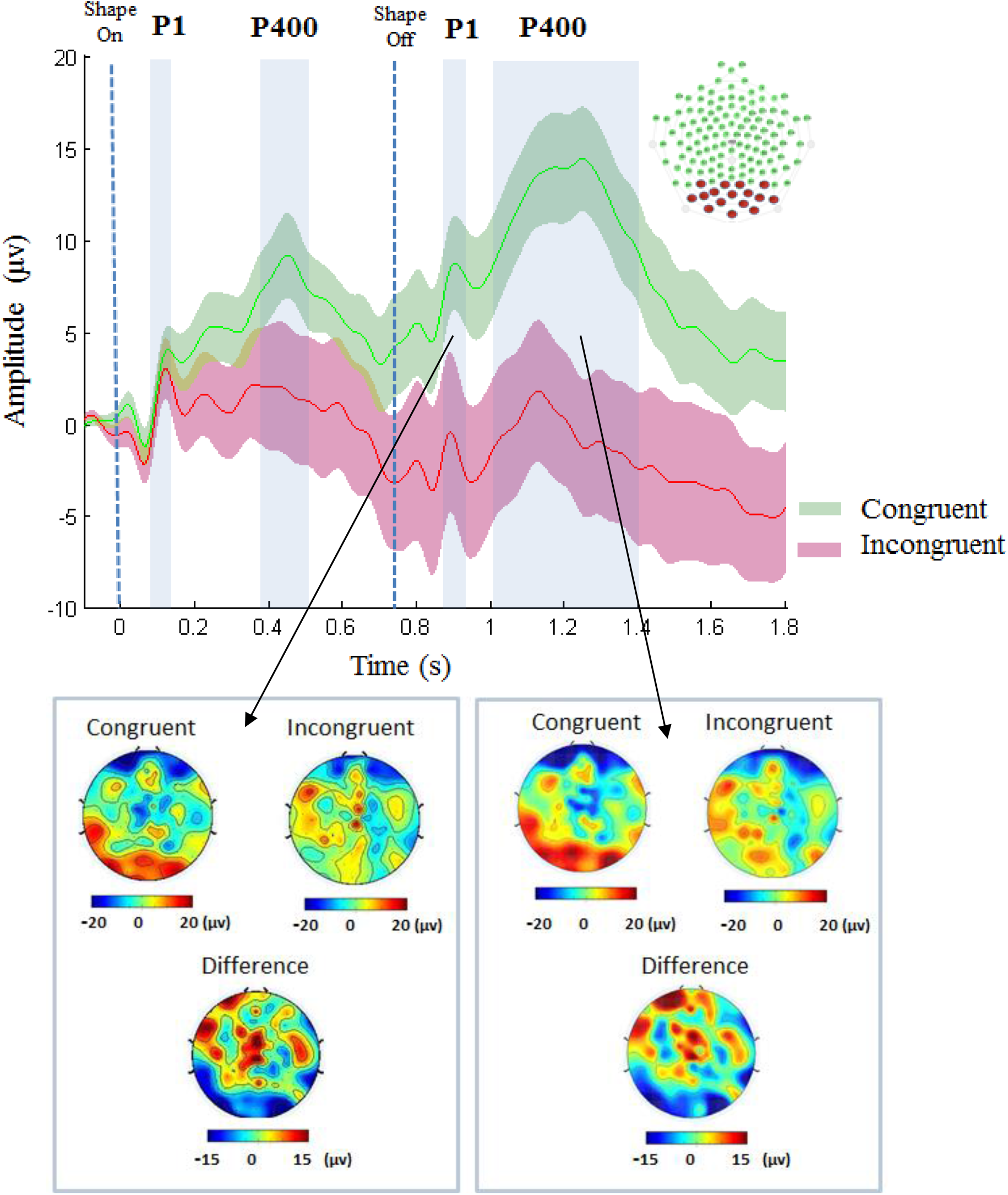
ERPs in response to congruent versus incongruent shapes and their topographies. **(A)** Grand-average in the congruent (green curve) and incongruent (red curve) conditions recorded on the occipito-temporal cluster of electrodes (red dots on the net topography). Blue rectangles indicate the time-window of the analysed ERP components. **(B)** Topographical maps of the grand average in each condition and their difference, during the P1 and P400 time-windows after the offset of the shape.

## Discussion

After a short training of 30 trials during which 3-month-old infants learned to associate two phonemes presented in different syllables with two specific shapes, infants displayed a difference in late ERPs for congruent and incongruent phoneme-shape pairings. Importantly, phonemes were presented in novel syllables different from those presented during familiarization, and which were furthermore paired with either of the two shapes with equal probability. As a result, no association between the novel syllables and the shapes could have been learned during the test phase. Instead, infants had to represent the abstract identity of the first consonant in order to discover the systematic association with the following visual shape.

As seen in figure 3, the difference between conditions slowly builds up from the time-window of the first P400 to significance only after the shape’s disappearance, corroborating the slowness of high-level processes frequently reported in infants (*13, 16*). This divergence culminated at the reappearance of the eyes. Moreover, the large response usually evoked by eyes in infants (*11, 12*) was significantly reduced in incongruent trials suggesting that infant’s attention was still focused on the previous erroneous shape pairing when the face reappeared. This prolonged effect is congruent with numerous studies reporting that infants have difficulties encoding new stimuli during stimulus orienting (*19, 20*). By contrast, we did not observe an early priming effect, characterised by larger early ERP components for the expected visual item, which has often been reported in audio-visual pairing tasks in older infants (*10, 17*). This might be due not only to the difficulty of the task but also the young age of our infants. The myelination of long-range tracts, necessary for fast transfer of information between brain regions (*21*) and efficient priming effects through top-down connections, is slow during the first year of life.

In our paradigm, infants were not directly discriminating consonants but were responding to a visual shape, which mediated the access to the consonants. Several experiments have reported that labelling helps infants categorize visual and auditory stimuli (*8–10*). A common arbitrary sign associated with a set of stimuli seems to encourage infants to search for shared features among stimuli, thereby directing their attention to the relevant features for analysis. The generalization to new syllables reveals that the shapes were not only an attention-attractor but that infants were using them somewhat like letters, that is, as explicit representations of the phonetic categories. Three-month-old infants can summarize a set of phonemic exemplars under the same visual label, well before they acquire their first words, revealing that phonemes are natural categories for pre-verbal infants.

Because phonemes are the smallest unit that can differentiate two words, it was hypothesized that infants begin to form phonetic representations once they notice that these small variations are relevant for distinguishing words (*22–25*). Our study refutes this hypothesis and is congruent with other observations suggesting that phonemes have an earlier perceptual relevance. 6-8 month-olds adapt their phonemic discrimination responses to the statistics of their environment (*26*), and 9 month-olds use contrastive pairings between objects and sounds to discover new phonemic categories (*27*). This can explain that already at 6-9 months, earlier than previously thought, infants understand a few words. Finally, it has been suggested that allophones (i.e., different realizations the same phoneme with no bearing on meaning) can be learned at a prelexical level (*28*) due to their complementary statistical distributions in words. This model proposed by Dupoux and Peperkamp implies that phonemes are pertinent units whose distributions can be computed regardless of their position in words and the surrounding phonemes. An interesting follow-up of our study would be to investigate the effect of position in the syllable (onset vs. coda) and whether infants would recognize that /ba/ and /ab/ share the same phoneme /b/.

In conclusion, our study shows that preverbal infants who can barely produce their first syllables and are only starting to vocalize can access sub-syllabic components and learn to map an initial consonant to an arbitrary visual shape in a few trials. Although our study was limited to two phonemes located at the onset of the syllable, this proof-of-concept reveals that the main cognitive process underlying reading, (i.e., creating a grapheme-phoneme pairing) is grounded in the natural faculties of the linguistic system.

## Materials and Method

### Participants

Thirty-three 3-month-old full-term infants raised in a French speaking environment were tested. Thirteen infants were not included in the analyses: 5 did not complete the experimental protocol, and 8 did not provide exploitable data due to poor data quality (see analysis section for exclusion criteria). The remaining 20 subjects comprised 8 girls and 12 boys, with a mean age of 14 weeks and 3 days (13 w. 5 d. to 16 w. 4 d.). The study was approved by the regional ethical committee for biomedical research, and the parents gave their written informed consent for the protocol.

### Auditory Stimuli

Ten syllables were naturally produced by a French female speaker (/gε/: 235 ms/, /gã/: 298 ms, 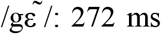, /gi/: 213 ms, /bε/: 244 ms, /bã/: 244 ms, and 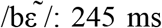 /bi/: 214 ms). We kept their initial duration to increase the variability of stimuli during familiarization. Two other syllables (/ba/: 199 ms and /ga/: 194 ms) produced by the same speaker and matched in duration were used as test stimuli. All syllables were matched for subjective intensity. Stimuli were recorded on the left channel and a click was positioned on the right channel at the exact time-point of syllable onset. The left channel was connected to an audio amplifier to monaurally present the sound to the subject while the right channel was connected to EEG amplifiers through the DIN port to send a TTL signal. Hence, when the sound was played by the PC soundcard, the brain voltage and the trigger signal were recorded simultaneously with the same temporal resolution.

### Visual Stimuli

Two still images of a woman were used, one with closed eyes and the second with open eyes. Only the upper part of the face (hair, forehead, and eyes) was presented against a blue background. These images were used as attentional fixation points for the infant. We also used two other images, a yellow cross and a red square tangent to a red rectangular bar as shapes to learn.

### Experimental Design

The female face remained on the screen during the entire experiment. Trials began with an eye blink 500 ms long. At the onset of the blink, a random syllable was presented (see Figure 1. for event duration). One second after the end of the blink, a shape appeared in the middle of the face and remained there for 766 ms. After a pause of random duration (ISI = 3200 to 3700 ms), the next trial began. The ISI jitter was used to increase the perception of pairs (sound-image) and avoid wrong associations between a shape and the following syllable. During familiarization, infants were first trained for 10 trials to one pairing (e.g. syllable /b/X and the yellow “+”), then to the other pairing for 10 trials (e.g. syllable /g/X followed by a red “%”), then both pairings were presented 5 times each in a random order. The order of presentation and phoneme-shape associations were counterbalanced across infants.

After the 30 trials of familiarization, the test phase began and the new syllables /ba/ and /ga/ were added to the previous set of syllables. These two syllables were followed half of the time by one shape and the other half by the other shape. The test comprised 20 short blocks of 8 randomly presented trials. Each one comprised four familiar trials (2 /b/X and 2 /g/X) to maintain learning and four new trials (2 /ba/ and 2 /ga/) followed with a shape either congruent or incongruent with those used during familiarization. No two incongruent trials followed one another. Short blocks were used to avoid imbalance between conditions in case the infant became agitated and the experiment terminated early.

### EEG recording and pre-processing

The 129-channel Geodesic sensor net (EGI) was placed on the infants’ heads relative to anatomical marks with the infant seated on their parents’ laps. Then, the infant and parent were seated facing a screen and a loudspeaker positioned behind the screen in a soundproof booth. The screen was located approximately 80 cm away from the infant’s face. If the infant looked away from the screen, the experiment was paused and the infant’s gaze was guided back to the screen before the experiment resumed. If this was not possible, the experiment was terminated.

Scalp voltages were referenced to the vertex, amplified (EGI, amplifiers N200), and digitized at 250 Hz. After recording, the raw data were filtered between 0.2 and 15 Hz and segmented into epochs starting 500 ms before and ending 1800 ms after the onset of the shape presentation. Artifact detection was done with custom-made scripts based on the EEGLAB toolbox (*29*). Epochs were considered unsuitable for analysis if their fast average amplitude exceeded 250 μV or their deviation between fast and slow running averages exceeding 150 μV. In each subject, channels that had more than 75% of epochs marked as bad were considered as bad channels. Five channels surrounding the eyes (17,125,126, 127 and 128) were rejected in several infants due to fixation difficulties. Therefore, we systematically rejected them in all subjects. For the remaining channels, 1.7 channels on average were rejected per infant. Trials having more than 50% of bad channels were rejected from the analysis. When a trial showed less than 25% of bad channels, these data were replaced by interpolating from neighbouring electrodes using spherical splines. After rejecting artifacted data, we obtained on average 81 acceptable trials per infant for the entire experiment with a mean number of rejected trials of 25.25 trials, and the number of good trials for the two conditions of interest (congruent/incongruent) was 13.7 and 13.6 (5 to 25 trials in each condition). Eight infants were rejected because they had less than 5 artefact-free trials in one of the two conditions. The retained epochs were re-referenced at each data-point to the mean voltage (*30*) to obtain a reference-free average and were finally baseline-corrected (−200 ms to −2 ms) in each subject.

### Statistical analysis

The analyses were performed on a set of 9 occipital and temporal electrodes comprising TP9, P9, PO7, O1, OZ, O2, PO8, P10 and TP10 (figure 2.), commonly used to study visual responses in infants (e.g. (*17*)). All trials were merged together to determine the time-windows for analysis, centered on P1 and P400. Voltage was averaged over electrodes and time-windows and entered 4 different t-test analyses (one for each component) with congruent vs. incongruent factors as a within-subject factor.

## References

1. G. Dehaene-Lambertz, M. Pena, Electrophysiological evidence for automatic phonetic processing in neonates. Neuroreport. 12, 3155–3158 (2001).

2. P. D. Eimas, E. R. Siqueland, P. Jusczyk, J. Vigorito, Speech perception in infants. Science. 171, 303–306 (1971).

3. J. Bertoncini, R. Bijeljac-Babic, P. W. Jusczyk, L. J. Kennedy, J. Mehler, An investigation of young infants’ perceptual representations of speech sounds. J. Exp. Psychol. Gen. 117, 21 (1988).

4. J.-R. Hochmann, L. Papeo, The invariance problem in infancy: A pupillometry study. Psychol. Sci. 25, 2038–2046 (2014).

5. K. Mersad, G. Dehaene-Lambertz, Electrophysiological evidence of phonetic normalization across coarticulation in infants. Dev. Sci. 19, 710–722 (2016).

6. J. Morais, L. Cary, J. Alegria, P. Bertelson, Does awareness of speech as a sequence of phones arise spontaneously? Cognition. 7, 323–331 (1979).

7. A. Karmiloff-Smith, Précis of Beyond modularity: A developmental perspective on cognitive science. Behav. Brain Sci. 17, 693–707 (1994).

8. A. L. Ferry, S. J. Hespos, S. R. Waxman, Nonhuman primate vocalizations support categorization in very young human infants. Proc. Natl. Acad. Sci. 110, 15231–15235 (2013).

9. H. H. Yeung, J. F. Werker, Learning words’ sounds before learning how words sound: 9-month-olds use distinct objects as cues to categorize speech information. Cognition. 113, 234–243 (2009).

10. Kabdebon C., G. Dehaene-Lambertz, Symbolic labelling in 5-month-old human infants. Submitted.

11. T. Gliga, G. Dehaene-Lambertz, Structural encoding of body and face in human infants and adults. J. Cogn. Neurosci. 17, 1328–1340 (2005).

12. T. Farroni, S. Massaccesi, D. Pividori, M. H. Johnson, Gaze following in newborns. Infancy. 5, 39–60 (2004).

13. G. Dehaene-Lambertz, S. Dehaene, Speed and cerebral correlates of syllable discrimination in infants. Nature. 370, 292 (1994).

14. G. Csibra, E. Kushnerenko, T. Grossmann, 15 Electrophysiological Methods in Studying Infant Cognitive Development. Handb. Dev. Cogn. Neurosci., 247 (2008).

15. A. Basirat, S. Dehaene, G. Dehaene-Lambertz, A hierarchy of cortical responses to sequence violations in three-month-old infants. Cognition. 132, 137–150 (2014).

16. S. Kouider et al., A neural marker of perceptual consciousness in infants. Science. 340, 376–380 (2013).

17. S. Kouider et al., Neural dynamics of prediction and surprise in infants. Nat. Commun. 6, 8537 (2015).

18. C. Kabdebon, G. Dehaene-Lambertz, Symbolic labelling in 5-month-old human infants. (Submitted).

19. J. E. Richards, Effects of attention on infants’ preference for briefly exposed visual stimuli in the paired-comparison recognition-memory paradigm. Dev. Psychol. 33, 22 (1997).

20. F. Leroy, R. Gulbinaite, L. Parkkonen, G. Dehaene-Lambertz, Attentional blink in infants: An electrophysiological and eye tracking study. Prep.

21. P. Adibpour, J. Dubois, G. Dehaene-Lambertz, How do electrophysiological measures in infants relate to the brain structural maturation? Neurophysiol. Clin. Neurophysiol. 47, 186 (2017).

22. C. T. Best, in Developmental neurocognition: Speech and face processing in the first year of life (Springer, 1993), pp. 289–304.

23. K. S. MacKain, Assessing the role of experience on infants’ speech discrimination. J. Child Lang. 9, 527–542 (1982).

24. J. F. Werker, R. C. Tees, Cross-language speech perception: Evidence for perceptual reorganization during the first year of life. Infant Behav. Dev. 7, 49–63 (1984).

25. J. F. Werker, R. C. Tees, Influences on infant speech processing: Toward a new synthesis. Annu. Rev. Psychol. 50, 509–535 (1999).

26. J. Maye, J. F. Werker, L. Gerken, Infant sensitivity to distributional information can affect phonetic discrimination. Cognition. 82, B101–B111 (2002).

27. H. H. Yeung, J. F. Werker, Learning words’ sounds before learning how words sound: 9-month-olds use distinct objects as cues to categorize speech information. Cognition. 113, 234–243 (2009).

28. E. Dupoux, S. Peperkamp, Fossil markers of language development: phonological ‘deafnesses’ in adult speech processing. Phon. Phonol. Cogn., 168–190 (2002).

29. A. Delorme, S. Makeig, EEGLAB: an open source toolbox for analysis of single-trial EEG dynamics including independent component analysis. J. Neurosci. Methods. 134, 9–21 (2004).

30. J. Dien, Issues in the application of the average reference: Review, critiques, and recommendations. Behav. Res. Methods Instrum. Comput. 30, 34–43 (1998).

